# Effects of hippocampal noninvasive theta-burst stimulation on consolidation of associative memory in healthy older adults

**DOI:** 10.1101/2023.10.11.554933

**Authors:** Traian Popa, Elena Beanato, Maximilian J. Wessel, Pauline Menoud, Fabienne Windel, Pierre Vassiliadis, Ines R. Violante, Ketevan Alania, Patrycja Dzialecka, Nir Grossman, Esra Neufeld, Friedhelm C. Hummel

## Abstract

Stimulation of deep brain areas can offer benefits against cognitive impairments associated with aging. So far, this was only possible via invasive methods accompanied by risks. Grossman *et al.* proposed a new noninvasive stimulation technique, transcranial temporal interference electric stimulation (tTIS), which can be steered to target and modulate activity of deep brain structures. Memory capacity depends on subcortical structures such as the hippocampus, hence, modulation of hippocampal activity could benefit declining cognitive functions. The current study investigates whether theta-burst patterned tTIS targeting the hippocampus influences performance of associative memory in older adults. We found that theta-burst patterned tTIS, but not the control stimulation, improved recollection time in a follow-up 24h after the stimulation, suggesting that theta-burst patterned tTIS can influence the efficiency of longer-term encoding. This outcome indicates that tTIS may provide a new noninvasive deep brain stimulation method to modulate senescent memory processes.

## Introduction

Associative memory has been shown to rely on medial temporal lobe and particularly, hippocampal activity^1–3^. Dysregulation or degeneration of this region lead to impaired memory functions that characterise aging^4–7^ and several neurological and neurodegenerative disorders^8,9^, such as Alzheimer’s disease^5,10,11^ or epilepsy^12,13^.

In healthy ageing in particular, a noticeable pattern of reduced ability to retain memories over longer time periods due to hippocampal changes14 and alterations in hippocampal-cortical and hippocampal-subcortical communication can be observed15–17.

Non-invasive brain stimulation (NIBS) techniques offer the possibility of modulating the activity of specific brain regions, resulting in behavioural effects^18^ that can be quantified and investigated. Focal, noninvasive stimulation of deep regions such as the hippocampus was until now not achievable without engagement of overlying cortical regions^19^, because of the steep depth-focality trade-off characterising traditional NIBS techniques^20^ (e.g., transcranial magnetic stimulation, TMS, or transcranial electrical stimulation, tES). The recently proposed tTIS method showed promising results for the noninvasive focal stimulation of deep regions in animals^21^. This technique combines two high-frequency currents that do not affect brain activity when applied independently, due to the dynamic properties of neurons (for details please see ^21,22^). The interference of the two electrical fields creates a low-frequency, beating envelope, which has been found, in contrast, to be able to modulate neural activity and whose modulation magnitude peak can be steered spatially inside the skull, even at depth^21^ (Fig. 2A). This technique has already shown promising results in humans when applied to both hippocampus^23^ and striatum^24,25^. Stimulation of the former led to increased accuracy in a face-name association task in healthy young participants. The same study also provided modelling and cadaver measures of the deep intracranial field supporting the good focality of the stimulation^23^. The striatum was successfully modulated as well, with tTIS leading to greater stimulation effects (motor learning capacity and fMRI striatal signal) especially in older with respect to young subjects^24^.

Theta-burst (tb-) stimulation pattern is a well-established protocol based on the characteristics of hippocampal physiology: complex-spike discharges of the pyramidal neurons^26^ and the rhythmic modulation of excitability of those cells during theta rhythms^27^. Combination of the complex-spike pattern (at 100Hz) with a theta frequency repetition rate of the bursts (at 5Hz) produces a robust, reliable, and stable LTP in the CA1 field of hippocampal slices^28^, which is more sensitive to many experimental conditions^29^.

In the current study, we investigated for the first time whether a theta burst pulsating field around the hippocampal circuits applied via tTIS (tb-TIS) would facilitate the intrinsic burst and subsequently enhance hippocampal memory processing in a cohort of healthy older adults^30–32^. Given that (i) the subparts of the hippocampus proper (i.e., *cornu amonis* and the dentate gyrus) are the most active during the encoding phase^33^, (ii) higher hippocampal activity during encoding is associated with better later recollection^33–35^, and (iii) direct hippocampal stimulation can enhance recollection^36,37^, we predicted better recollection for face-name pairs encoded during tb-tTIS.

## Results

To investigate the effects of hippocampal tb-tTIS on memory performance, we employed a well-established face-name association task^23,38^. Both accuracy and response times of correct associations were extracted from the main and follow-up trials (Fig. 1). The participants were asked to respond as accurately and as fast as possible when primed to choose the correct name for the faces displayed during the retrieval part of each trial. The participant’s confidence in the accuracy of the response given was assessed for each answer on a scale 0-5 (“unsure” to “absolutely sure”).

**Fig 1.**
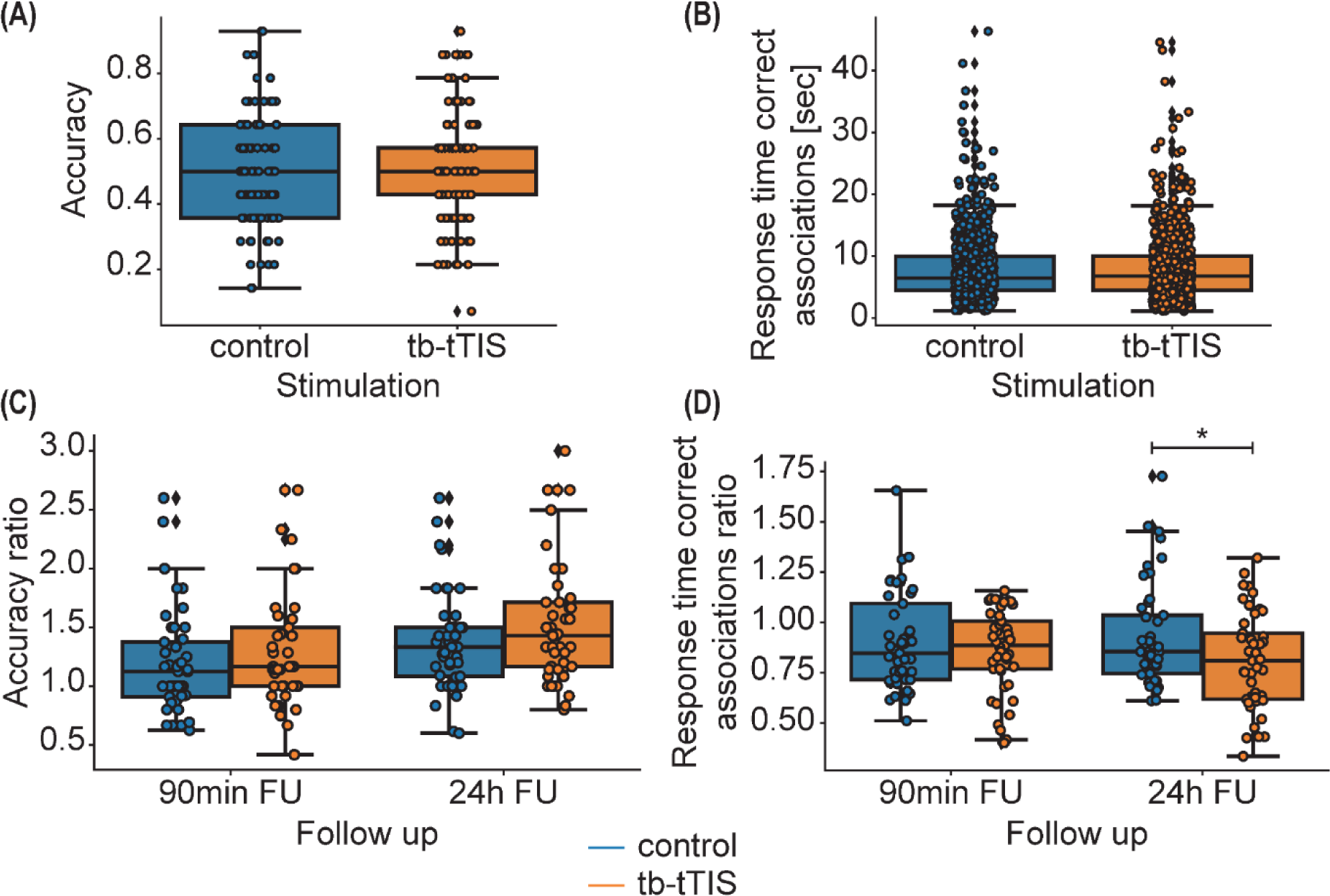
Behavioural results. (A) Distribution of the accuracy during stimulation blocks. No significant difference was found. (B) Distribution of the trial response time of correct associations during stimulation blocks. No significant difference was found. (C) Accuracy during the follow-up sessions 90 minutes and 24 hours after stimulation, with respect to the last block of the main session. No significant difference was found. (D) Average response time for the correct associations over each block of the follow-up sessions 90 minutes and 24 hours after stimulation, with respect to the average response time for the correct associations during the last block of stimulation. A significant difference was found between stimulation conditions, with hippocampal tb-tTIS leading to faster retrieval of the correct associations (*t(18.2) = -2.23, p = 0.04, d = -0.94 [large]*).

When running a generalised linear mixed model on the accuracy during the stimulated blocks, ‘stimulation’ was used as a fixed factor and ‘subject’ as a random intercept. The distribution was assumed to be binomial by using the *logit* link function. There was no significant effect of stimulation (*X*^2^*(1, 15) = 0.02, p = 0.88, d = 0.01 [micro]*). The effect of the stimulation on response times of correct associations during the main session was investigated via a linear mixed model on log-transformed data (due to the data distribution). The winning model included ‘stimulation’ as a fixed effect, ‘subject’ and ‘block’ as random intercepts, and ‘stimulation’ as the random slope for the factor ‘subject’. There was no significant effect of the stimulation (*F(1,14.02) = 0.03, p = 0.87, pη*^2^ *= 0.002 [micro]*). These results indicate that tb-tTIS did not induce changes in memory retrieval right after the first encoding of the information. To investigate potential recall effects, we first tested the difference in performance during the last block with stimulation between the two stimulation conditions. No difference was found in accuracy or response time of correct associations (accuracy: paired Wilcoxon test, *V=44.5, p=0.64, d=-0.07; Bayesian paired t-test, BF10 = 0.27, moderate evidence*; response times: paired Wilcoxon test, *V=85, p=0.17, d=-0.4; Bayesian paired t-test, BF10 = 0.7, anecdotal evidence*). The performance of the last block was hence used as reference for the follow-ups. The level of performance change during follow-ups with respect to the main session was quantified as the average performance divided by the performance during the last stimulated block and statistically assessed. The winning model included again ‘stimulation’ and ‘follow-up’ as fixed factors, ‘subject’ as random intercept, and ‘stimulation’ as random slope. Results indicated a significant stimulation x follow-up interaction (*F(1, 148) = 4.74, p = 0.03, pη*^2^*= 0.03 [small]*). This was driven by faster responses during the 24 hour, but not the 90 minute, follow-up after hippocampal tb-tTIS compared to the control stimulation (*t(18.2) = -2.23, p = 0.04, d = -0.94 [large]*). These results support the ability of hippocampal tb-tTIS to influence associative memory, specifically the time to retrieve correct answers probed during the consolidation period. A linear mixed model with ‘stimulation’ and ‘follow-up’ as fixed factors, ‘subject’ as random intercept, and ‘stimulation’ as random slope, revealed no statistically significant improvement in accuracy after tb-tTIS with respect to control stimulation (*F(1, 14) = 0.65, p = 0.43, pη*^2^ *= 0.04 [small]*). The confidence associated with correct associations was also analysed both during main and follow-up sessions as a secondary outcome. No stimulation effect was found (for more details, see section “Confidence” in SOM).

## Discussion

The majority of the studies inducing behavioural effects on associative memory in humans were obtained via either direct, invasive or indirect, noninvasive cortical stimulation approaches. Only one recent study provided evidence that direct, noninvasive tTIS^23^ in young healthy subjects is able to modulate associative memory. Studies targeting the hippocampus or adjacent regions of the medial temporal lobe with invasive deep brain stimulation (DBS) in human neurosurgical cases typically used a 50 Hz stimulation frequency^39–44^. This approach led to a variety of contrasting results and different research groups reported either disrupting^39,41^, beneficial^40,43,44^, or no effects on memory^42^. We found only one study applying theta-burst patterned stimulation in humans, similar to the theta-burst protocol used in the current study, via invasive microstimulation of the hippocampal afferent inputs^45^. The theta-burst patterned invasive stimulation induced positive effects on memory recall that are in line with the present results, although those improvements were observed already during the first recall (the response time was not reported, and there was no follow-up probing). The divergence in behavioural stimulation outcomes of previous studies could be explained by several factors. Firstly, they used different memory tasks, which could engage different hippocampal territories, and hence lead to different stimulation effects, as suggested by Jun and colleagues (2020)^44^. Secondly, duration and intensity of the stimulation^43^, as well as the location and the focality^45^, could each significantly influence the outcome. Despite these divergences, all studies conclude that the hippocampus is crucially involved in declarative memory and that it is possible to modulate behaviour via hippocampal stimulation.

Because of the limitation in reaching the hippocampus noninvasively, another branch of studies focused on the modulation of hippocampal activity with indirect non-invasive cortical stimulation methods. More precisely, theta-burst transcranial magnetic stimulation was used on cortical areas linked to the hippocampus, with the aim of indirectly affecting hippocampal activity via cortico-hippocampal connections. These studies showed that indirect entrainment of the hippocampal activity is possible, and it can lead to physiological and behavioural changes^46–49^. It is worth noting that these positive effects were obtained when specifically targeting the encoding period^50^, also in line with the currently employed protocol.

Finally, the study targeting the left hippocampus of young healthy participants with direct, noninvasive tTIS found higher retrieval accuracy during tTIS with respect to no stimulation^23^. The discrepancy between previous and current behavioural effects of tTIS could be explained by multiple differences in the experimental protocol: (1) Violante and colleagues employed a continuous, not bursted, theta tTIS, suggested to entrain ongoing theta hippocampal activity, in contrast to the present theta-burst pattern that is suggested to induce LTP-like phenomena in the hippocampus; (2) the stimulation was applied throughout both encoding and retrieval of a face-name association task, in contrast to the encoding-only stimulation applied here; (3) the peak of the envelope modulation was steered toward the anterior part of the hippocampus, in contrast to the center of the hippocampus targeted in the current study; and (4) most importantly, results were extracted from a young healthy cohort, in contrast to the older cohort in our experiments. These previous results demonstrate that tTIS can achieve focal modulation of hippocampal activity; the comparison with our outcomes suggests that variations in the protocol might differently impact memory mechanisms.

In the current study, significant stimulation effects were specifically found during the 24 hours follow-up, but not during the main session. This opens the discussion for interesting insights about the *modus operandi* of tb-tTIS stimulation within the hippocampus. The major difference between the main and follow-up sessions is that all face-name associations were novel during the stimulated blocks, while they were re-encoded during both follow-ups, *i.e.*, at 90min and 24h post-stimulation (*N.B.,* each main session had its own set of face-name associations). It was hypothesised that second and third presentations may benefit later memory by providing an opportunity for extended processing of the name^51^. While it is expected that this would lead to higher accuracy and lower response time during the repetition of already seen associations^52–54^, tb-tTIS boosted these effects even further. Previous fMRI studies in healthy, young adults revealed that repeated presentations of the same face-name associations lower hippocampal activation and reduce the neocortical deactivations in the antero-medial and postero-medial areas^51,55,56^. These patterns were found altered in cognitively healthy older individuals^57,58^ and patients with Alzheimer’s disease^59^, who were specifically characterised by a failure of task-associated deactivation of cortical regions during encoding. The continuous interaction between the hippocampus and its ample connected cortical network^60^ is critical for successful episodic memory encoding and retrieval, requiring a coordinated reciprocal pattern of activations and deactivations in response to repeated stimuli^55^. Although we could not perform simultaneous fMRI recordings to quantify the variations in hippocampal and neocortical activations and in their functional connectivity at each time-point, we can speculate about possible modes of action of hippocampal tb-tTIS. One possibility is that tb-tTIS successfully entrained the natural hippocampal theta bursts, supporting hippocampal encoding activity. By facilitating the initial encoding of face-name pairs, subsequent presentations could reinforce and enhance the association of already optimised pairs^56^. This process could improve accessibility of the stored information, ultimately leading to faster recollection. Another possibility is that tb-tTIS improved the coordination of the theta-bursts characterising multimodal associative encoding^61^. This could subsequently lead to a more efficient communication from the hippocampus to other networks involved in this process, in line with a network effect as suggested by the indirect noninvasive stimulation results. Increased theta content in hippocampal activity could hence lead to more efficient cortical deactivations, which have been proven to be important for subsequent retrievals^62^. Finally, we cannot exclude that both of these mechanisms could play a role in the observed behavioural effects. This last hypothesis is further supported by the observation that decision time depends on cortical encoding strength, which in turn depends on encoding hippocampal activity^63^. Furthermore, cortical reinstatement (i.e., the reactivation of specific cortical patterns representing previously encoded information) during retrieval is related to the cortical activity during encoding^63^, which could explain the faster response time observed in the current study.

The presence of effects during the 24 hours follow-up only was unexpected. This discovery remains to be investigated with further behavioural and targeted neuroimaging studies. The hypothesis we had about tTIS driving the behavioural improvements via either local or network effects would not explain the observed dichotomy between the 90 minutes and the 24 hours follow-up effects. However, a possible explanation could be related to a specific impact of the stimulation on the later consolidation processes occurring between the two follow-up sessions, namely during sleep^64^. The benefits of sleep for long-term memory consolidation are well established^64^. During this process, specific oscillatory patterns are hypothesised to enhance communication between the hippocampus and cortical areas to redistribute memory traces^65^. Certain memories were shown to be preferentially consolidated based on their characteristics^66^ and the level of the encoding^67,68^, poorly encoded memories being prioritised for consolidation during sleep^66^. These considerations could be especially relevant if the participants present inefficient memory consolidation^69^ due to age-related structural and functional alterations^70^. In line with this hypothesis, age-related impairments were demonstrated to particularly emerge in poorly encoded memories before sleep^71^, whilst no difference was found with respect to young subjects for strong memory representations. By boosting the encoding strength of the face-name pairs, hippocampal tb-tTIS might reduce the need for consolidation mechanisms that are impaired in an older cohort, hence leading to the positive behavioural effects observed here.

Some considerations about the novel tTIS technique should be highlighted to better interpret the findings. Modelling studies of tTIS effects suggested that in humans, the electric fields created by intensities tolerable for the scalp (in our case, maximum 2mA / channel) would be strong enough to entrain intrinsic oscillations or indirectly modulate spiking activity, but not strong enough to trigger action potentials within deep brain structures^72–74^. Hence, the observed behavioural changes would be in line with sub-threshold neuromodulation effects. However, assumptions based on traditional NIBS techniques need to be considered with caution and further investigated. In fact, it is unclear how a given low-frequency tTIS envelope would impact brain activity in comparison to a unique, low-frequency alternating field, and indeed if the modulation of the amplitude is the most relevant exposure quantity-of-interest. Hence, the magnitude of the fields required to induce neuromodulation could differ between techniques. It is worth noting that HF exposure can result in reduced sensations and increased tolerability^73,75^, which would theoretically allow exploration of higher stimulation intensities.

Taken together, these results provide first evidence that the hippocampus can be directly noninvasively neuromodulated using the innovative tTIS technology in older adults with a consequent impact on associative memory. Theta-burst patterned tTIS demonstrated the ability to induce faster recall of previously encoded face-name pairs 24 hours after stimulation when compared to a control stimulation. This evidence sets the ground for further investigations on the role of subcortical regions in memory functions, which is fundamental for a better understanding of both healthy as well as pathological human systems. tTIS could be used as an important tool to explore how ageing affects hippocampal functioning and how this is linked to well-characterised age-related cognitive impairments. Importantly, the present study highlights the potential of tTIS as a neuromodulation tool for future translation into neurorehabilitation and therapeutic applications.

## Methods

### Participants

Fifteen healthy older adults (right-handed, 9 female, average age 66.07 ± 5.57 years old) were recruited for the current project. The detailed exclusion criteria are listed in SOM. The study was conducted in accordance with the Declaration of Helsinki and approved by the Cantonal Ethics Committee Vaud, Switzerland (project number 2020-00127). All participants gave their written informed consent.

### Study design

#### Cognitive task

Episodic memory ability was investigated with a face-name association task^23,38^ (Fig. 2C). The task was separated into alternating encoding and retrieval blocks. Each encoding block consisted of a series of 14 faces, each presented with a unique name written underneath. The faces were displayed one after the other, for 3 seconds each. Each retrieval block consisted of the same 14 faces, each represented with five names underneath: the correct name, two names used during the encoding but associated with other faces (foil names), and two names never seen during the encoding (distractor names). The participants were asked to select the name they remembered to be associated with the face and, after each choice, to give an estimation of their confidence level on a scale from 1 to 4 (with 1 corresponding to low confidence and 4 corresponding to high confidence). The duration of the retrieval block hence depended on the response time of each subject. All participants were instructed to perform the task “as fast and as accurate as possible”.

**Fig 2.**
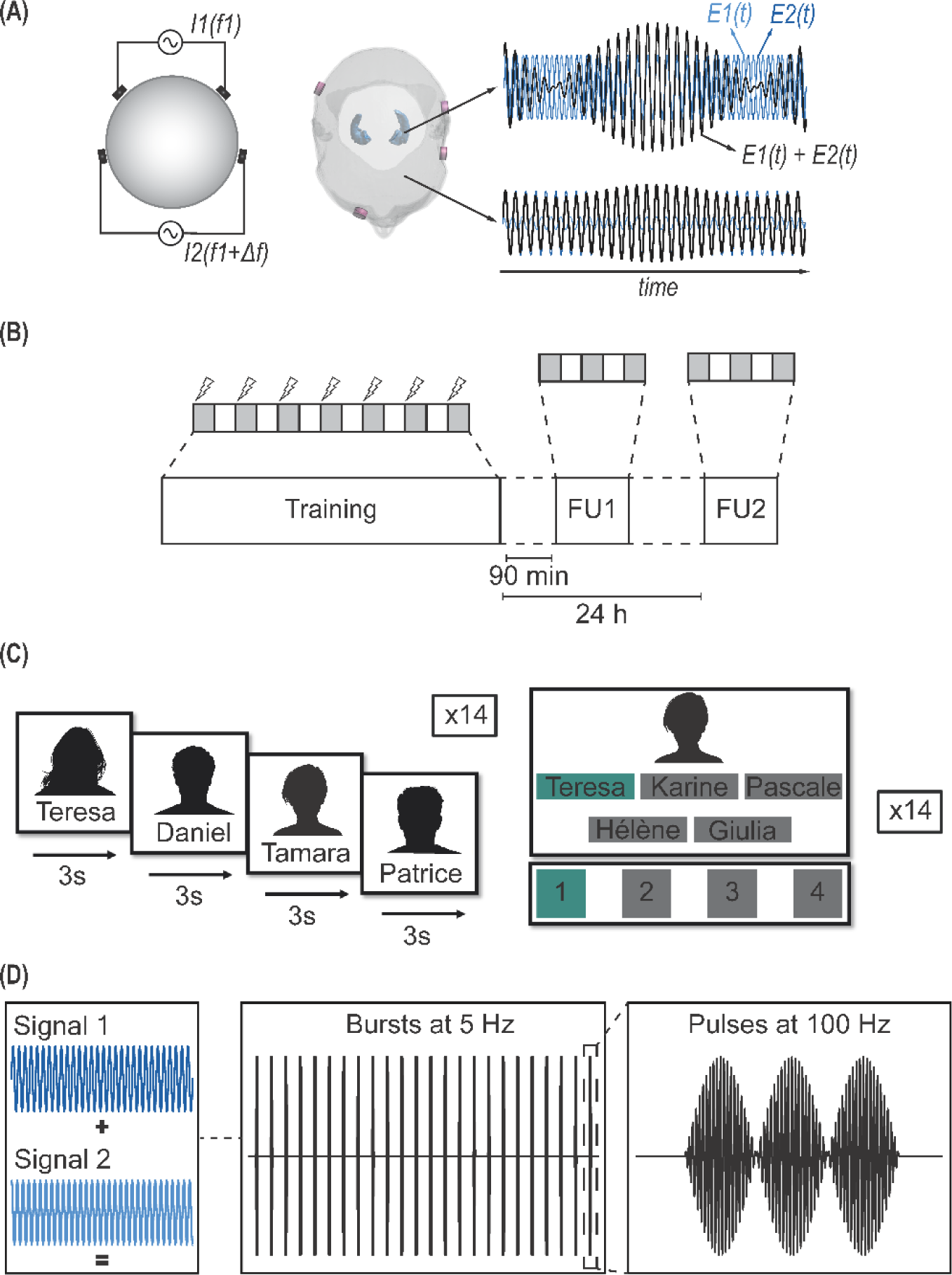
Protocol design and stimulation patterns. (A) Transcranial Temporal Interference Stimulation concept. From left to right: spherical head model with two pairs of electrodes delivering currents at a frequency f1 and f1+Δf; real head model with hippocampus masks highlighted in blue; the resulting interference of the two electric fields is conceptually illustrated for a point in the target structure and for a remote one. The envelope modulation is evident within the target region and weak outside (plots are illustrative and were not extracted from simulations or recordings). (B) Protocol of the study. During the main session, participants performed seven blocks of a face-name association task, with concomitant stimulation during the encoding period only. They were then asked to perform three of the seven blocks 90 minutes and 24 hours later. (C) Face-name association task. During the first part of the task (encoding), 14 faces were shown one after another, for 3 seconds each, along with an associated name. During the second part (retrieval), each of the faces were shown again with 5 possible names underneath. Subjects had to choose the name they remember to be associated with the face and, once selected, give an estimate of their confidence on a scale from 1 (not confident at all) to 4 (sure). (D) Theta-burst protocol created by the interference of the two high-frequency currents. The two signals are combined to form bursts at a frequency of 5 Hz, each composed of three pulses at 100 Hz. In the periods between pulses or bursts, the current was not switched off, but the two carrier frequencies were equalised. The control high-frequency stimulation consisted only in two unmodulated, equal carrier frequencies.

#### Experimental protocol

The study had a randomised, double-blind, cross-over design, with each subject undergoing a total of five visits. During the first one, after the signature of the informed consent, safety questionnaires and baseline evaluations were performed. The safety questionnaires included the check of the inclusion criteria for transcranial stimulations and the magnetic resonance environment screening form. The baseline evaluations included demographic data, current work situation and educational level, list of ongoing and past medications, medical history, handedness (Edinburgh Laterality Test^76^), sleep quality (Pittsburg Sleep Quality Index, PSQI^77^), Frontal Assessment Battery^78^ (FAB) and MOntreal Cognitive Assessment^79^ (MOCA).

The second visit started by performing seven consecutive blocks of the face-name association task. The participants received either the active or the control stimulation during the encoding phase of each block. Three of the seven blocks were then repeated without any stimulation 90 minutes after the end of the seven blocks with stimulation and 24 hours after (the third visit). The fourth and fifth visits were identical to the second and third (main session, 90 minutes and 24 hours follow-up), but with the other stimulation protocol. The two stimulation visits (third and fifth) were performed with at least one week of interval between them, to minimise any potential carryover effects.

The Stanford Sleepiness Scale80 (SSS) and three Visual Analog Scales (VAS) to assess fatigue were administered prior to the beginning of the main and follow-up sessions, and the VAS were reassessed right after each of them.

#### Stimulation protocol

All subjects received both an active and an HF control stimulation protocol, both involving two HF current delivery channels^21,81^. When the two channel frequencies differ slightly, their superposition features a modulation envelope oscillating at their difference frequency. However, when that frequency shift is reset, no such modulation occurs, resulting in a constant high-frequency exposure. By applying specific frequency shifts at specific timings, an active and a control stimulation protocol were created. The active stimulation mimicked a theta burst stimulation pattern applied continuously: bursts of three pulses at 100 Hz were delivered every 200 ms (5 Hz). The control stimulation consisted of a flat envelope, with no shift in frequency between the two current channels. Currents were ramped up in the beginning of each block of the task and ramped down at the end of the encoding phase. The following parameters were used: carrier frequency = 2.0 kHz (resp. 2.1 kHz for one channel in the active condition), current intensity per stimulation channel = 2 mA, duration = 7 min (including ramp-up and ramp-down), ramp-up = 10 sec, ramp-down = 8 sec, electrode type: round, conductive rubber with conductive cream/paste, electrode size = 3 cm^2^.

### Stimulation

#### Placement of the electrodes and stimulation-associated sensations

The location of the stimulating electrodes was taken from Violante *et al.*^23^ in order to target the left hippocampus (please see Fig. 2C). The electrode pairs were hence placed around the P8-CP7 and Fp2-FT7 landmarks, respectively, in the 10-20 EEG electrode positions system^82^. Locations were marked with a pen on the skin. After skin preparation (cleaned with alcohol), the round conductive rubber electrodes were placed, adding a conductive paste (Ten20, Weaver and Company, Aurora, CO, USA or Abralyt HiCl, Easycap GmbH, Woerthsee-Etterschlag, Germany) as interface with the skin, and held in position with tape. The electrodes were placed with cables protruding downwards, to minimise movements due to gravity, except for the electrode on the front, which was tilted horizontally in order not to occlude the visual field of participants. Impedances were optimised until they were lower than 20 kΩ. Once good contacts were obtained, the subjects received different current intensities of short duration to familiarise them with the perceived sensations and to systematically document their feedback: both the tb-tTIS and control stimulation were applied for 20 seconds in separate rounds, with increasing current amplitudes from 0.5 mA, 1 mA, 1.5 mA, up to 2 mA per channel. Participants were asked to report any kind of sensation and for each intensity, if any sensation was felt, they were asked to grade its strength from 1 to 3 (*i.e.,* light to strong) and give at least one adjective to describe it. A nonconductive, elastic cap was then used to apply slight pressure on the electrodes and keep them in place. Additional impedance checks were performed immediately before and after the stimulation to check the maintenance of a good contact.

### Cognitive task analysis

Two main variables were extracted from the face-name association task: accuracy and speed. During the main session, accuracy was taken as a binomial variable 0 or 1 (not correct or correct association) for each trial. Response time was measured as the time between the appearance of the face and the selection of the name. Only response times of correct associations were considered. During the retrieval phase, accuracy was computed as the percentage of correct associations per block, whilst response time of correct associations was averaged per block. Both variables were then corrected by dividing by the respective performance during the last block of stimulation. The corrected average confidence was also calculated in line with previously described methods, as a secondary outcome (please see SOM).

### Statistical analysis

The statistical analysis was performed in R software (R Core Team (2021). R: A language and environment for statistical computing. R Foundation for Statistical Computing, Vienna, Austria, https://www.R-project.org/; version 4.0.5). Depending on the distribution of the data, generalised or simple linear mixed models (*glmer* or *lmer* functions from the *lme4* package^83^) were used to test the differences during main and follow-up sessions. For the accuracy during the main session, the binomial family was used, whilst for the other models, variables were checked for normality (Shapiro-Wilk test) and if data were not normally distributed, the log transform was applied. For the main session results, different models were hierarchically implemented. They included ‘‘stimulation’’ as a fixed factor, adding step by step ‘subject’ and ‘block’ as random intercepts, and ‘stimulation’’ as a random slope for the ‘subject’ factor. For the retention data, different models were hierarchically implemented including ‘stimulation’ and ‘follow-up’ (90min or 24h after) as fixed factors, ‘subject’ as random intercept, and ‘stimulation’, which was added as random slope when possible. In both cases, the best model was chosen via likelihood tests implemented with the *anova* function (from the *stats* package^84^). The *anova* function with Satterthwaite’s approximations (from the *lmerTest* package^85^) was then run on the final model. Post-hoc analyses were also conducted via pairwise comparisons by computing estimated marginal means with the *emmeans* package^86^. For questionnaire outcomes, paired t-test or Wilcoxon rank sum test were used, based on the normality of the data, which was tested with the Shapiro-Wilk test. All functions were provided by the *stats* package^84^. Finally, effect size measures were obtained using the *effect size* package^87^ and are expressed as partial eta-squared^88^ (pη^2^, pη^2^: < 0.01 ∼ micro, 0.01 ∼ small, 0.06 ∼ medium, 0.14 ∼ large) for F-tests and Cohen’s d^88^ (d: < 0.2 ∼ micro, 0.2-0.3 ∼ small, 0.5 ∼ medium, 0.8 ∼ large effect size) for paired comparisons. The level of significance was set at p < 0.05.

Because of the low sample size, we also reported Bayesian statistics in the SOM obtained with the *brms* package^89,90^. The uninformative default priors were used because of the lack of a previous hypothesis on the tTIS effects. The same model as the one chosen via the likelihood test of the frequentist models was specified. Posterior probabilities were then used to compute the probability of observing the effect under investigation.

## Acknowledgements

We would like to thank A.G. Cadic-Melchior, P. Maceira and T. Morishita for their assistance with the randomization, and the neuromodulation facility of the Human Neuroscience Platform of the Fondation Campus Biotech Geneva for technical advice.

## Data availability

The data and scripts necessary to generate the main results are available in the Zenodo repository.

---

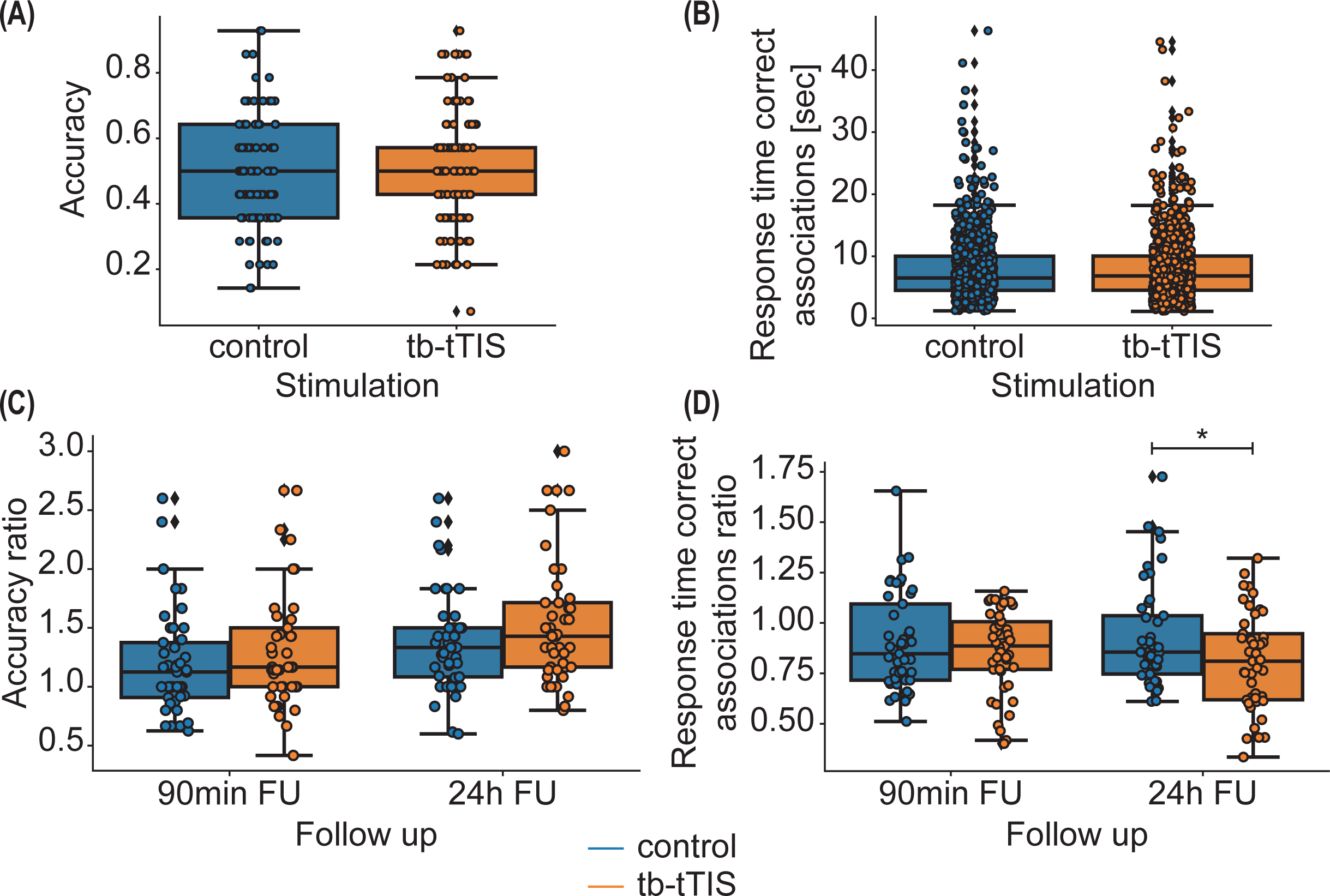

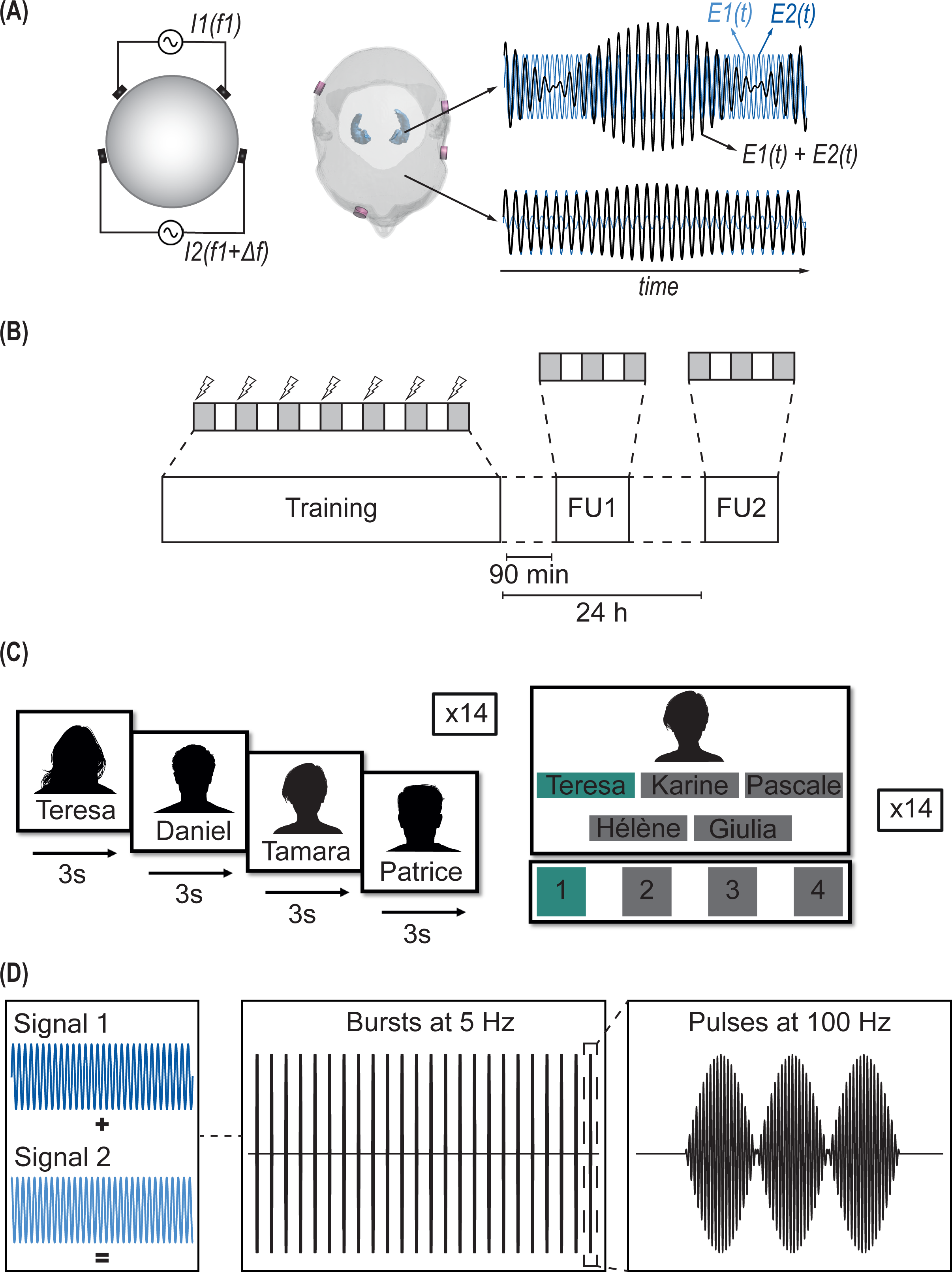

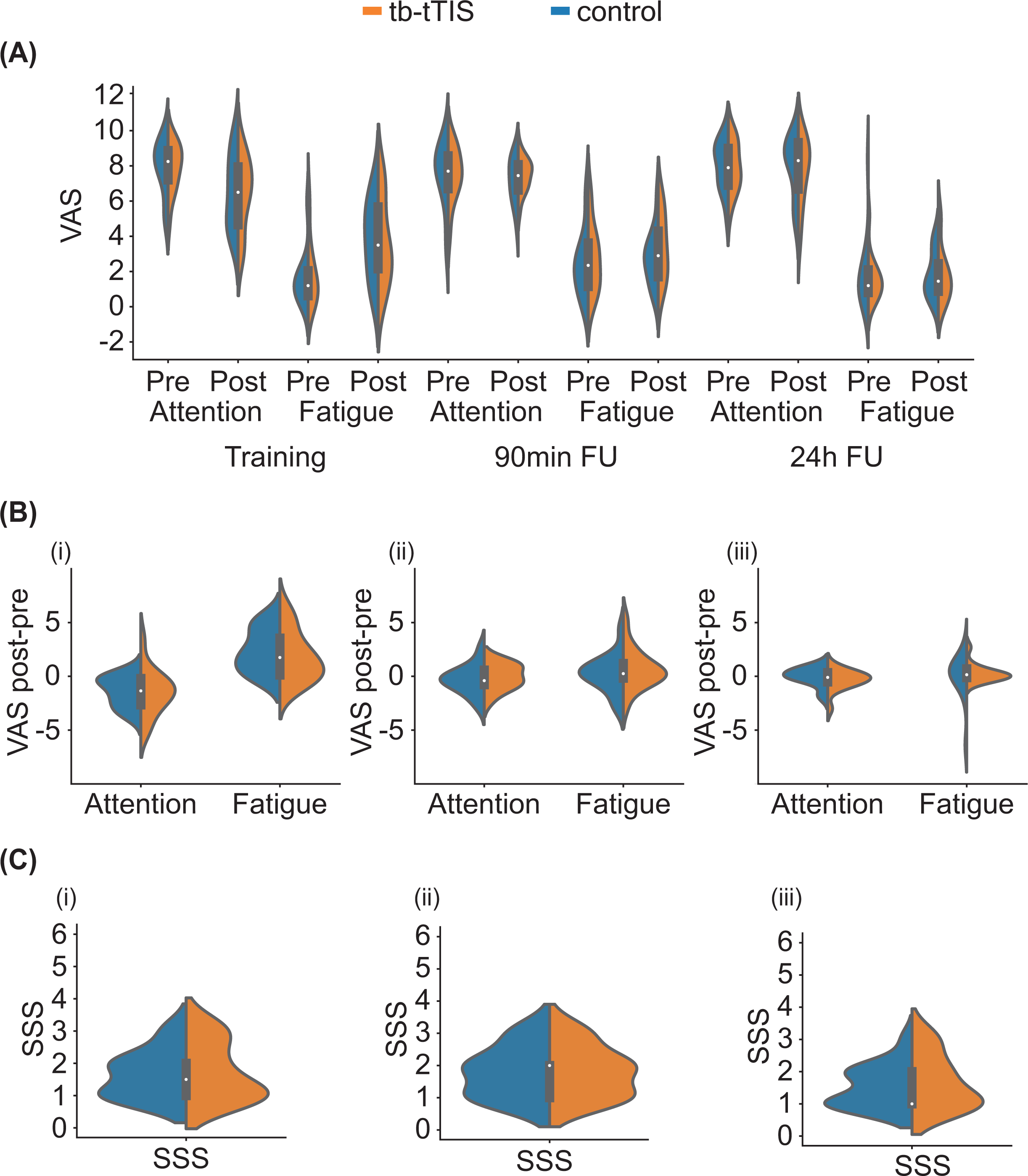

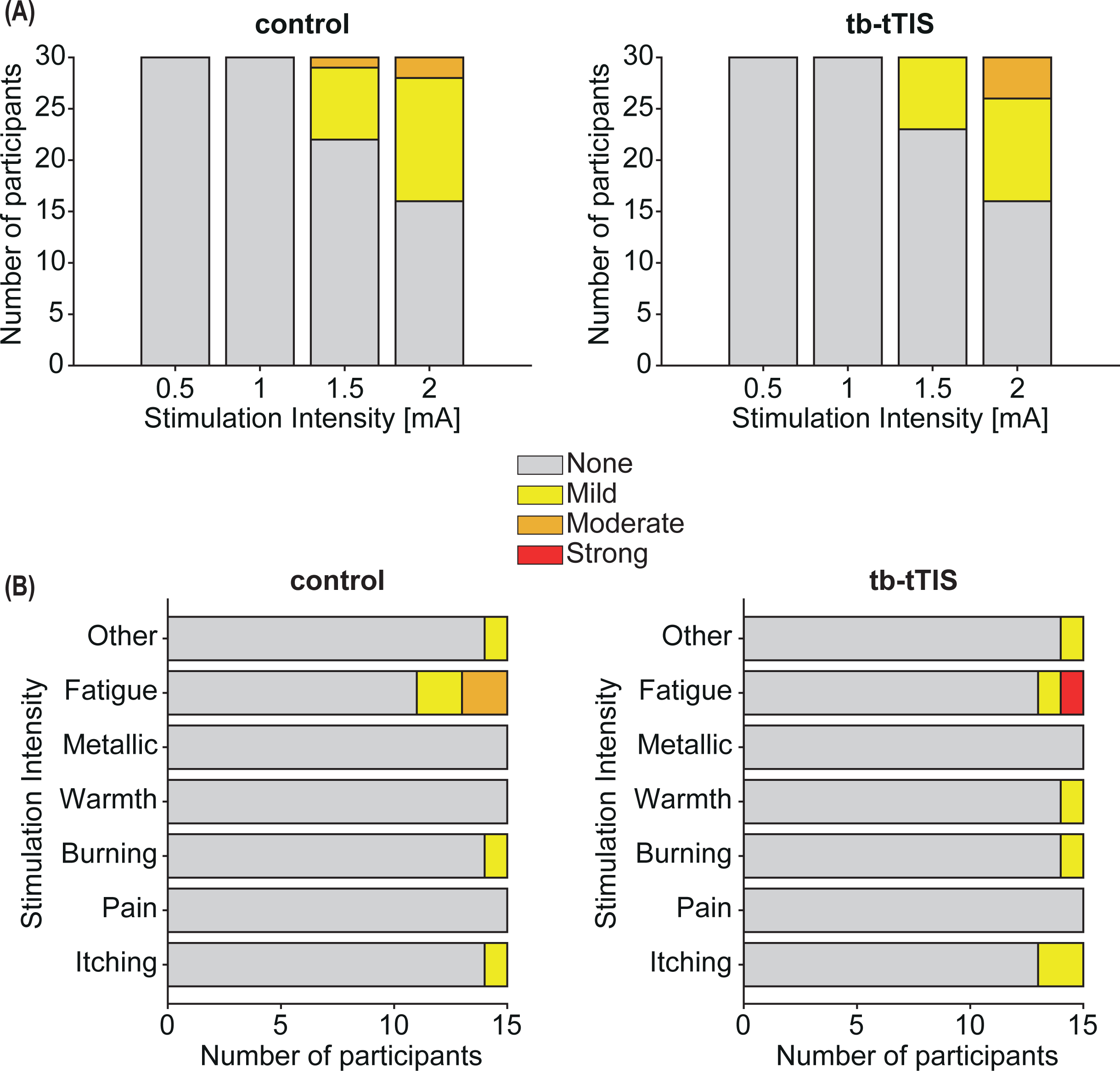

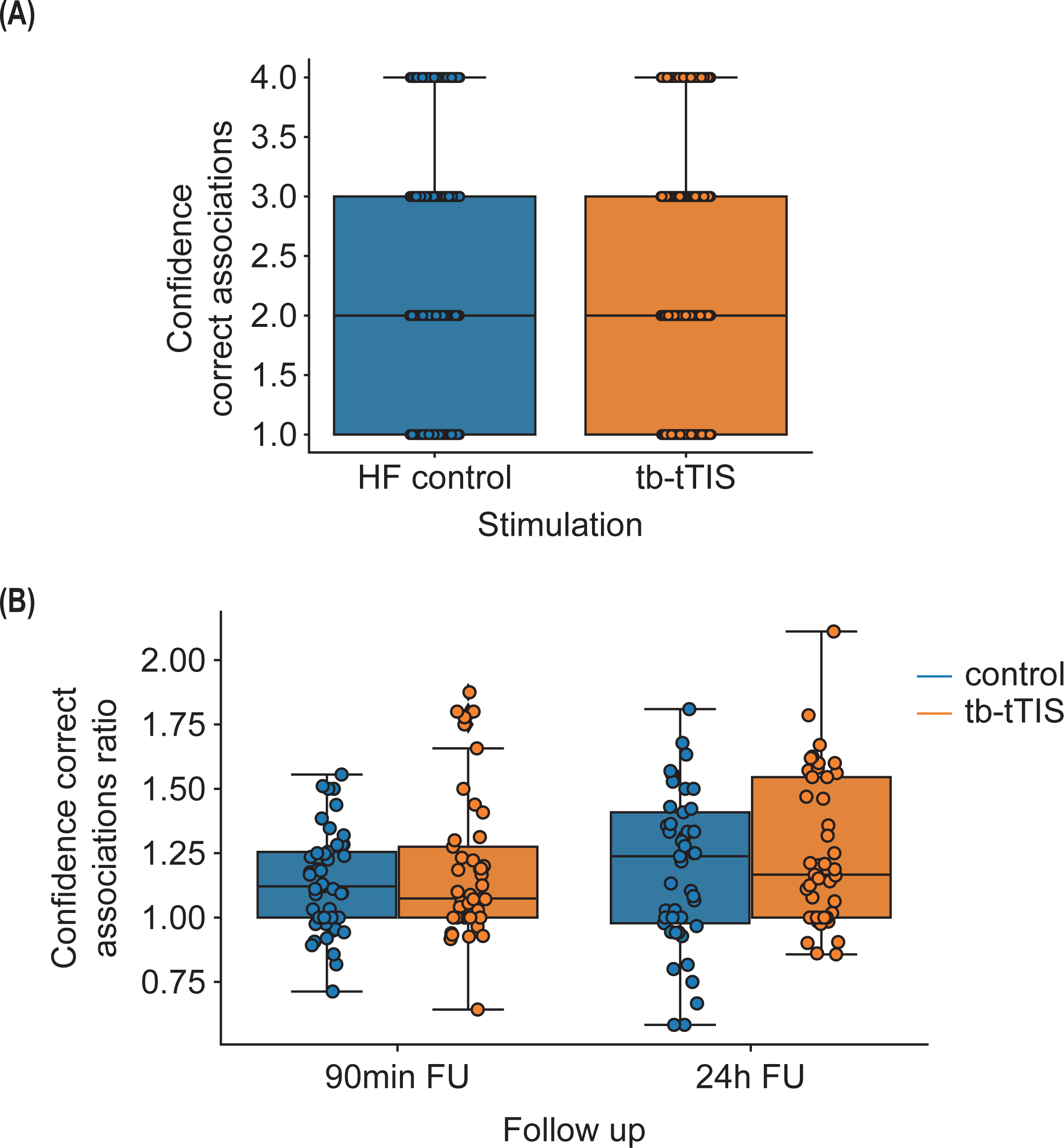

## Bibliography

1. Scoville, W. B. & Milner, B. Loss of recent memory after bilateral hippocampal lesions. J. Neurol. Neurosurg. Psychiatry 20, 11–21 (1957).

2. Squire, L. R. Declarative and nondeclarative memory: multiple brain systems supporting learning and memory. J. Cogn. Neurosci. 4, 232–243 (1992).

3. Opitz, B. Memory function and the hippocampus. Front. Neurol. Neurosci. 34, 51–59 (2014).

4. Lister, J. P. & Barnes, C. A. Neurobiological changes in the hippocampus during normative aging. Arch. Neurol. 66, 829–833 (2009).

5. Tromp, D., Dufour, A., Lithfous, S., Pebayle, T. & Després, O. Episodic memory in normal aging and Alzheimer disease: Insights from imaging and behavioral studies. Ageing Res. Rev. 24, 232–262 (2015).

6. Wheeler, M. A. A comparison of forgetting rates in older and younger adults. Aging Neuropsychol. Cogn. 7, 179–193 (2000).

7. MacDonald, S. W., Stigsdotter-Neely, A., Derwinger, A. & Bäckman, L. Rate of acquisition, adult age, and basic cognitive abilities predict forgetting: new views on a classic problem. J. Exp. Psychol. Gen. 135, 368 (2006).

8. Moodley, K. K. & Chan, D. The hippocampus in neurodegenerative disease. Front. Neurol. Neurosci. 34, 95–108 (2014).

9. Weerasinghe-Mudiyanselage, P. D. E., Ang, M. J., Kang, S., Kim, J.-S. & Moon, C. Structural Plasticity of the Hippocampus in Neurodegenerative Diseases. Int. J. Mol. Sci. 23, 3349 (2022).

10. Braak, H. & Braak, E. [Morphology of Alzheimer disease]. Fortschr. Med. 108, 621–624 (1990).

11. Mu, Y. & Gage, F. H. Adult hippocampal neurogenesis and its role in Alzheimer’s disease. Mol. Neurodegener. 6, 85 (2011).

12. Hoppe, C., Elger, C. E. & Helmstaedter, C. Long-term memory impairment in patients with focal epilepsy. Epilepsia 48 Suppl 9, 26–29 (2007).

13. Fleury, M. et al. Episodic memory network connectivity in temporal lobe epilepsy. Epilepsia 63, 2597–2622 (2022).

14. Eichenbaum, H. Hippocampus: cognitive processes and neural representations that underlie declarative memory. Neuron 44, 109–120 (2004).

15. Leshikar, E. D., Gutchess, A. H., Hebrank, A. C., Sutton, B. P. & Park, D. C. The impact of increased relational encoding demands on frontal and hippocampal function in older adults. Cortex J. Devoted Study Nerv. Syst. Behav. 46, 507–521 (2010).

16. Salami, A., Pudas, S. & Nyberg, L. Elevated hippocampal resting-state connectivity underlies deficient neurocognitive function in aging. Proc. Natl. Acad. Sci. U. S. A. 111, 17654–17659 (2014).

17. Ness, H. T. et al. Reduced Hippocampal-Striatal Interactions during Formation of Durable Episodic Memories in Aging. Cereb. Cortex N. Y. N 1991 32, 2358–2372 (2022).

18. Koch, G. & Caltagirone, C. Non-invasive brain stimulation: From brain physiology to clinical opportunity. Neurosci. Lett. 719, 134496 (2020).

19. Lee, S., Lee, C., Park, J. & Im, C.-H. Individually customized transcranial temporal interference stimulation for focused modulation of deep brain structures: a simulation study with different head models. Sci. Rep. 10, 11730 (2020).

20. Deng, Z.-D., Lisanby, S. H. & Peterchev, A. V. Electric field depth-focality tradeoff in transcranial magnetic stimulation: simulation comparison of 50 coil designs. Brain Stimulat. 6, 1–13 (2013).

21. Grossman, N. et al. Noninvasive Deep Brain Stimulation via Temporally Interfering Electric Fields. Cell 169, 1029–1041.e16 (2017).

22. Hutcheon, B. & Yarom, Y. Resonance, oscillation and the intrinsic frequency preferences of neurons. Trends Neurosci. 23, 216–222 (2000).

23. Violante, I. R., et al. Non-invasive temporal interference electrical stimulation of the human hippocampus. http://biorxiv.org/lookup/doi/10.1101/2022.09.14.507625(2022) doi:10.1101/2022.09.14.507625.

24. Wessel, M. J., et al. LTP-like noninvasive striatal brain stimulation enhances striatal activity and motor skill learning in humans. http://biorxiv.org/lookup/doi/10.1101/2022.10.28.514204(2022) doi:10.1101/2022.10.28.514204.

25. Vassiliadis, P., et al. Non-invasive stimulation of the human striatum disrupts reinforcement learning of motor skills. http://biorxiv.org/lookup/doi/10.1101/2022.11.07.515477(2022) doi:10.1101/2022.11.07.515477.

26. Ranck, J. B. Studies on single neurons in dorsal hippocampal formation and septum in unrestrained rats. I. Behavioral correlates and firing repertoires. Exp. Neurol. 41, 461– 531 (1973).

27. Rudell, A. P., Fox, S. E. & Ranck, J. B. Hippocampal excitability phase-locked to the theta rhythm in walking rats. Exp. Neurol. 68, 87–96 (1980).

28. Larson, J., Wong, D. & Lynch, G. Patterned stimulation at the theta frequency is optimal for the induction of hippocampal long-term potentiation. Brain Res. 368, 347–350 (1986).

29. Larson, J. & Munkácsy, E. Theta-burst LTP. Brain Res. 1621, 38–50 (2015).

30. Lisman, J. E. & Jensen, O. The theta-gamma neural code. Neuron 77, 1002–1016 (2013).

31. Hanslmayr, S., Axmacher, N. & Inman, C. S. Modulating Human Memory via Entrainment of Brain Oscillations. Trends Neurosci. 42, 485–499 (2019).

32. Rudoler, J. H., Herweg, N. A. & Kahana, M. J. Hippocampal Theta and Episodic Memory. J. Neurosci. Off. J. Soc. Neurosci. 43, 613–620 (2023).

33. Zeineh, M. M., Engel, S. A., Thompson, P. M. & Bookheimer, S. Y. Dynamics of the hippocampus during encoding and retrieval of face-name pairs. Science 299, 577–580 (2003).

34. Paller, K. A. & Wagner, A. D. Observing the transformation of experience into memory. Trends Cogn. Sci. 6, 93–102 (2002).

35. Jun, S., Kim, J. S. & Chung, C. K. Hippocampal Neuronal Activity Preceding Stimulus Predicts Later Memory Success. eNeuro 10, ENEURO.0252-22.2023 (2023).

36. Hebscher, M. & Voss, J. L. Testing network properties of episodic memory using non-invasive brain stimulation. Curr. Opin. Behav. Sci. 32, 35–42 (2020).

37. Kota, S., Rugg, M. D. & Lega, B. C. Hippocampal Theta Oscillations Support Successful Associative Memory Formation. J. Neurosci. Off. J. Soc. Neurosci. 40, 9507–9518 (2020).

38. Sperling, R. A. et al. Putting names to faces: successful encoding of associative memories activates the anterior hippocampal formation. NeuroImage 20, 1400–1410 (2003).

39. Coleshill, S. G. et al. Material-specific recognition memory deficits elicited by unilateral hippocampal electrical stimulation. J. Neurosci. Off. J. Soc. Neurosci. 24, 1612–1616 (2004).

40. Suthana, N. A. et al. Memory enhancement and deep-brain stimulation of the entorhinal area. N. Engl. J. Med. 366, 502–510 (2012).

41. Jacobs, J. et al. Direct Electrical Stimulation of the Human Entorhinal Region and Hippocampus Impairs Memory. Neuron 92, 983–990 (2016).

42. Hansen, N. et al. Memory encoding-related anterior hippocampal potentials are modulated by deep brain stimulation of the entorhinal area. Hippocampus 28, 12–17 (2018).

43. Jun, S., Kim, J. S. & Chung, C. K. Direct Stimulation of Human Hippocampus During Verbal Associative Encoding Enhances Subsequent Memory Recollection. Front. Hum. Neurosci. 13, 23 (2019).

44. Jun, S., Lee, S. A., Kim, J. S., Jeong, W. & Chung, C. K. Task-dependent effects of intracranial hippocampal stimulation on human memory and hippocampal theta power. Brain Stimulat. 13, 603–613 (2020).

45. Titiz, A. S. et al. Theta-burst microstimulation in the human entorhinal area improves memory specificity. eLife 6, (2017).

46. Hermiller, M. S., Chen, Y. F., Parrish, T. B. & Voss, J. L. Evidence for Immediate Enhancement of Hippocampal Memory Encoding by Network-Targeted Theta-Burst Stimulation during Concurrent fMRI. J. Neurosci. 40, 7155–7168 (2020).

47. Hermiller, M. S. et al. Evidence from theta-burst stimulation that age-related de-differentiation of the hippocampal network is functional for episodic memory. Neurobiol. Aging 109, 145–157 (2022).

48. Thakral, P. P., Madore, K. P., Kalinowski, S. E. & Schacter, D. L. Modulation of hippocampal brain networks produces changes in episodic simulation and divergent thinking. Proc. Natl. Acad. Sci. 117, 12729–12740 (2020).

49. Hermiller, M. S., VanHaerents, S., Raij, T. & Voss, J. L. Frequency-specific noninvasive modulation of memory retrieval and its relationship with hippocampal network connectivity. Hippocampus 29, 595–609 (2019).

50. Tambini, A., Nee, D. E. & D’Esposito, M. Hippocampal-targeted Theta-burst Stimulation Enhances Associative Memory Formation. J. Cogn. Neurosci. 30, 1452–1472 (2018).

51. Mangels, J. A., Manzi, A. & Summerfield, C. The First Does the Work, But the Third Time’s the Charm: The Effects of Massed Repetition on Episodic Encoding of Multimodal Face–Name Associations. J. Cogn. Neurosci. 22, 457–473 (2010).

52. Stevenage, S. V. & Spreadbury, J. H. Haven’t we met before? The effect of facial familiarity on repetition priming. Br. J. Psychol. 97, 79–94 (2006).

53. Johnston, R. A. & Barry, C. Repetition priming of access to biographical information from faces. Q. J. Exp. Psychol. 59, 326–339 (2006).

54. Takashima, A. et al. Memory trace stabilization leads to large-scale changes in the retrieval network: a functional MRI study on associative memory. Learn. Mem. Cold Spring Harb. N 14, 472–479 (2007).

55. Vannini, P., Hedden, T., Sullivan, C. & Sperling, R. A. Differential functional response in the posteromedial cortices and hippocampus to stimulus repetition during successful memory encoding. Hum. Brain Mapp. 34, 1568–1578 (2013).

56. Rand-Giovannetti, E. et al. Hippocampal and neocortical activation during repetitive encoding in older persons. Neurobiol. Aging 27, 173–182 (2006).

57. Miller, S. L. et al. Age-related memory impairment associated with loss of parietal deactivation but preserved hippocampal activation. Proc. Natl. Acad. Sci. 105, 2181– 2186 (2008).

58. Sperling, R. A. et al. Amyloid Deposition Is Associated with Impaired Default Network Function in Older Persons without Dementia. Neuron 63, 178–188 (2009).

59. Pihlajamäki, M., O’Keefe, K., O’Brien, J., Blacker, D. & Sperling, R. A. Failure of repetition suppression and memory encoding in aging and Alzheimer’s disease. Brain Imaging Behav. 5, 36–44 (2011).

60. Robinson, J. L., Salibi, N. & Deshpande, G. Functional connectivity of the left and right hippocampi: Evidence for functional lateralization along the long-axis using meta-analytic approaches and ultra-high field functional neuroimaging. NeuroImage 135, 64–78 (2016).

61. Battaglia, F. P., Benchenane, K., Sirota, A., Pennartz, C. M. A. & Wiener, S. I. The hippocampus: hub of brain network communication for memory. Trends Cogn. Sci. 15, 310–318 (2011).

62. Kaefer, K., Stella, F., McNaughton, B. L. & Battaglia, F. P. Replay, the default mode network and the cascaded memory systems model. Nat. Rev. Neurosci. 23, 628–640 (2022).

63. Gordon, A. M., Rissman, J., Kiani, R. & Wagner, A. D. Cortical reinstatement mediates the relationship between content-specific encoding activity and subsequent recollection decisions. Cereb. Cortex N. Y. N 1991 24, 3350–3364 (2014).

64. Rasch, B. & Born, J. About sleep’s role in memory. Physiol. Rev. 93, 681–766 (2013).

65. Cowan, E. et al. Sleep Spindles Promote the Restructuring of Memory Representations in Ventromedial Prefrontal Cortex through Enhanced Hippocampal-Cortical Functional Connectivity. J. Neurosci. Off. J. Soc. Neurosci. 40, 1909–1919 (2020).

66. Brodt, S., Inostroza, M., Niethard, N. & Born, J. Sleep-A brain-state serving systems memory consolidation. Neuron 111, 1050–1075 (2023).

67. Denis, D. et al. Sleep Spindles Preferentially Consolidate Weakly Encoded Memories. J. Neurosci. Off. J. Soc. Neurosci. 41, 4088–4099 (2021).

68. Zhang, J., Whitehurst, L. N. & Mednick, S. C. The role of sleep for episodic memory consolidation: Stabilizing or rescuing? Neurobiol. Learn. Mem. 191, 107621 (2022).

69. Faßbender, R. V. et al. Decreased Efficiency of Between-Network Dynamics During Early Memory Consolidation With Aging. Front. Aging Neurosci. 14, 780630 (2022).

70. Muehlroth, B. E., Rasch, B. & Werkle-Bergner, M. Episodic memory consolidation during sleep in healthy aging. Sleep Med. Rev. 52, 101304 (2020).

71. Muehlroth, B. E. et al. Memory quality modulates the effect of aging on memory consolidation during sleep: Reduced maintenance but intact gain. Neuroimage 209, 116490 (2020).

72. Rampersad, S., et al. Prospects for transcranial temporal interference stimulation in humans: a computational study. http://biorxiv.org/lookup/doi/10.1101/602102(2019) doi:10.1101/602102.

73. Esmaeilpour, Z., Kronberg, G., Reato, D., Parra, L. C. & Bikson, M. Temporal interference stimulation targets deep brain regions by modulating neural oscillations. Brain Stimulat. 14, 55–65 (2021).

74. Howell, B. & McIntyre, C. C. Feasibility of Interferential and Pulsed Transcranial Electrical Stimulation for Neuromodulation at the Human Scale. Neuromodulation J. Int. Neuromodulation Soc. 24, 843–853 (2021).

75. Hsu, G., Farahani, F. & Parra, L. C. Cutaneous sensation of electrical stimulation waveforms. Brain Stimulat. 14, 693–702 (2021).

76. Oldfield, R. C. The assessment and analysis of handedness: the Edinburgh inventory. Neuropsychologia 9, 97–113 (1971).

77. Smyth, C. The Pittsburgh Sleep Quality Index (PSQI). J. Gerontol. Nurs. 25, 10–10 (1999).

78. Dubois, B., Slachevsky, A., Litvan, I. & Pillon, B. The FAB: A frontal assessment battery at bedside. Neurology 55, 1621–1626 (2000).

79. Nasreddine, Z. S. et al. Montreal Cognitive Assessment. (2014) doi:10.1037/t27279-000.

80. Hoddes, E., Zarcone, V., Smythe, H., Phillips, R. & Dement, W. C. Quantification of sleepiness: a new approach. Psychophysiology 10, 431–436 (1973).

81. Grossman, N. Modulation without surgical intervention. Science 361, 461–462 (2018).

82. Seeck, M. et al. The standardized EEG electrode array of the IFCN. Clin. Neurophysiol. Off. J. Int. Fed. Clin. Neurophysiol. 128, 2070–2077 (2017).

83. Bates, D., Mächler, M., Bolker, B. & Walker, S. Fitting Linear Mixed-Effects Models using lme4. (2014) doi:10.48550/ARXIV.1406.5823.

84. Bolar, K. STAT: interactive document for working with basic statistical analysis. (2019).

85. Luke, S. G. Evaluating significance in linear mixed-effects models in R. Behav. Res. Methods 49, 1494–1502 (2017).

86. Lenth, R. & Lenth, M. R. Package ‘lsmeans’. Am. Stat. 34, 216–221 (2018).

87. Ben-Shachar, M. S., Lüdecke, D. & Makowski, D. effectsize: Estimation of effect size indices and standardized parameters. J. Open Source Softw. 5, 2815 (2020).

88. Cohen, J. Statistical power analysis. Curr. Dir. Psychol. Sci. 1, 98–101 (1992).

89. Bürkner, P.-C. brms : An *R* Package for Bayesian Multilevel Models Using *Stan*. J. Stat. Softw. 80, (2017).

90. Bürkner, P.-C. Advanced Bayesian Multilevel Modeling with the R Package brms. R J. 10, 395 (2018).

