## Supplementary material for "Effects of hippocampal noninvasive theta-burst stimulation on consolidation of associative memory in healthy older adults"

### Baseline questionnaires

Participants were characterised at baseline via several questionnaires summarised in the following table:

| Sample size | Age | EHI | FAB | MOCA | PSQI |
|---|---|---|---|---|---|
| 15 | 66.07±5.57 | 90.32±14.18 | 15.80±2.48 | 26.33±1.99 | 4.73±2.05 |

Attention and fatigue were measured before and after each main and follow up session (Fig. S1A-B). No effect of stimulation was detected (pre-post main session attention: $t(14)=-0.21$, $p=0.84$, $d=-0.06$; pre-post main session fatigue: $V=31$, $p=0.33$, $d=-0.27$; pre-post FU1 attention: $t(14)=0.81$, $p=0.43$, $d=0.22$; pre-post FU1 fatigue: $V=66.5$, $p=0.73$, $d=0.24$; pre-post FU2 attention: $t(12)=-0.21$, $p=0.84$, $d=-0.07$; pre-post FU2 fatigue: $V=37.5$, $p=0.60$, $d=0.09$). Statistics for pre-post comparison of both fatigue and attention during FU2 are computed on 13 subjects because of missing data. Sleepiness was measured before starting the task during main session and follow ups (Fig. S1C). No significant differences were found between sessions with active or control stimulation (before main session: $V=25$, $p=0.80$, $d=0.08$; before FU1: $V=18.5$, $p=1.00$, $d=0$; before FU2: $V=26.5$, $p=0.67$, $d=0.18$).

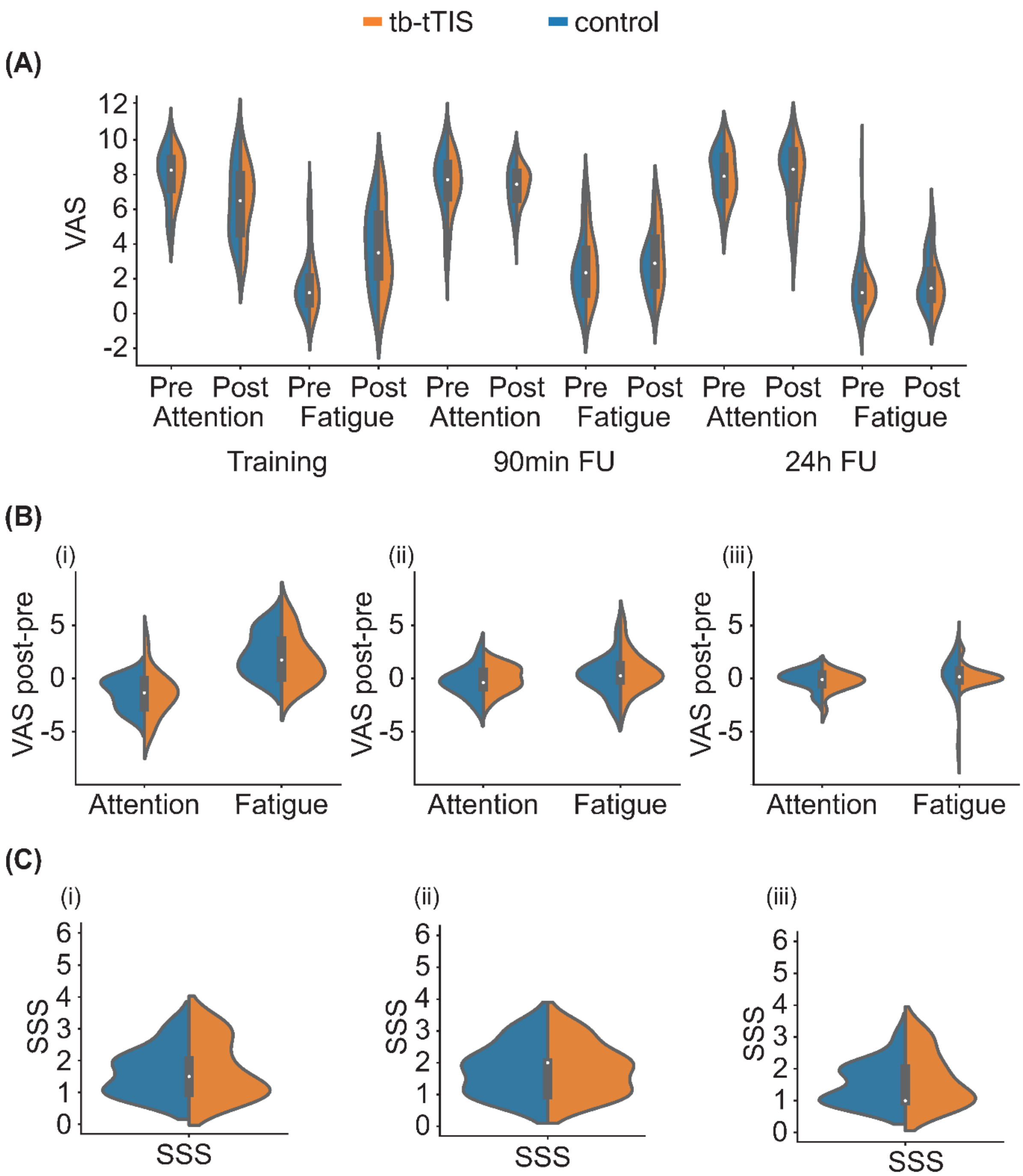


**Fig S1. Fatigue and attention**
(A) Attention and fatigue levels measured via the Visual Analog Scales (VAS), before and after main and follow-up sessions (90 minutes and 24 hours after). (B) Task-induced changes in attention and fatigue levels. No significant changes were observed. (i) Changes from before to after the main session. (ii) Changes from before to after the 90 minutes follow up. (iii) Changes from before to after the 24 hours follow-up. (C) Stanford Sleepiness Scale (SSS) before (i) main session, (ii) 90 minute follow-up and (iii) 24 hours follow-up. No significant changes were observed.

### tTIS associated sensation and blinding

A successful blinding of the study was confirmed by multiple tests (Fig. S2). Firstly, the level of tTIS-associated sensations during the initial testing was comparable between tb-tTIS and control stimulation, as confirmed by the non-significant stimulation x intensity factor (*F(3,203)=0.29, p=0.84,* $p\eta^2$*=0.004 [micro]*). Secondly, reported sensations during the task also revealed a non-significant difference between the two stimulation conditions (*Z=-0.514, p=0.607*). Finally, when asked to guess which stimulation protocol they received, participants' responses did not deviate from chance-level guessing (exact binomial test: *p* = *1*).

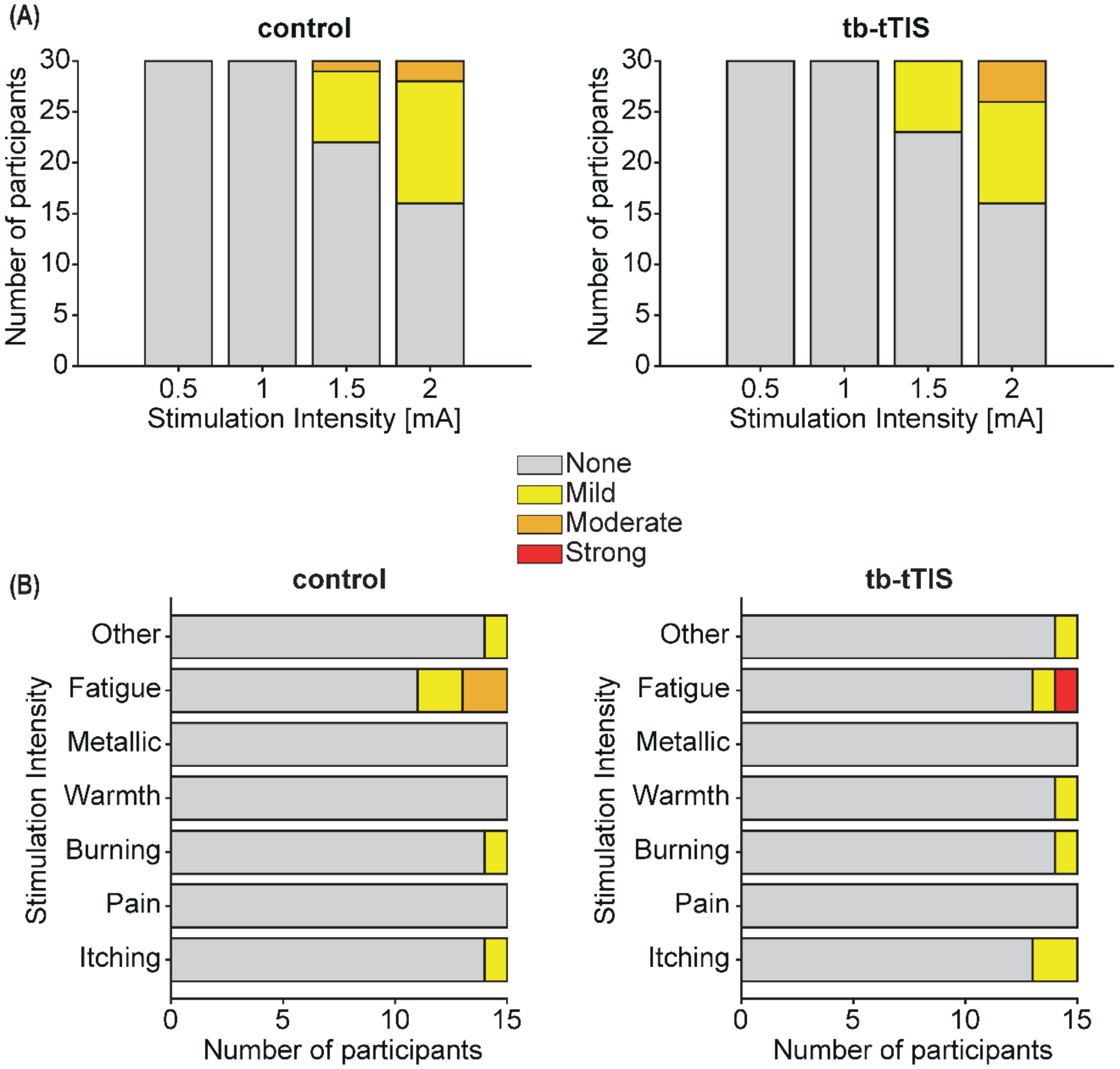


**Fig S2. tTIS induced sensations**

(A) Sensation perceived during the sensation test. Subjects received around 20 seconds of each stimulation protocol, from a current intensity of 0.5 to 2 mA, increasing with steps of 0.5 mA. The graphs report the sensation felt during control stimulation, on the left, and tb-tTIS, on the right. No significant difference in stimulation perception was found (stimulation x condition interaction: *F(3,203)=0.29, p=0.84,* $\eta p^2$*=0.004 [micro]*). (B) Sensations perceived during the association task with concomitant stimulation. Subjects were asked at the end of the task to describe which sensations they felt. Results are plotted for control stimulation on the left and for tb-tTIS on the right. No significant effect of the stimulation condition was found (*Z=-0.514, p=0.607*).

#### Bayesian statistics

In this section we report Bayesian statistics supporting the current findings. All results are described as the probability of observing a difference between the stimulation conditions, with 50% representing random chances. During stimulating blocks, there was a probability of 55.48% and 55.63% of observing an increased accuracy and reaction time respectively, during tb-tTIS with respect to control stimulation. This is in line with the absence of a significant stimulation effect. During the follow ups, there was a probability of 97.71% to observe faster response times 24 hours after tb-tTIS than after control stimulation, whilst 77.68% probability of observing higher accuracy. Again, this confirms the large effect of tb-tTIS on the response time.

#### Confidence

We additionally investigated whether the stimulation affected confidence levels of correct associations during main or follow up sessions. During the main session, because of the nature of the data including integer values from one to four, we used a generalised linear mixed model with a Poisson distribution and a logarithmic link function. The winning model included stimulation as fixed factor and subject as random intercept. No significant effect of stimulation was observed on the confidence level (*X2 (1, 15) = 2.76, p = 0.10, d = -0.06 [micro]*), even though Bayesian statistics resulted in a probability of 95.14% of observing a higher confidence during control stimulation with respect to tb-tTIS.

During the retrieval period, data were corrected over the last block similarly to the analysis performed on accuracy and response time. Data was not normally distributed even after logarithmic transformation, but close to normal as verified by the *descdist* function in R. We therefore implemented a linear mixed model with stimulation and follow up as fixed factors, subject as random intercept and stimulation as random slope. No effect of the stimulation was present ($F(1, 14) = 0.07$, $p = 0.79$, $p\eta^2 = 0.005$ *[micro]*), with 60.19% probability of observing higher confidence after tb-tTIS with respect to control (50% being chance level).

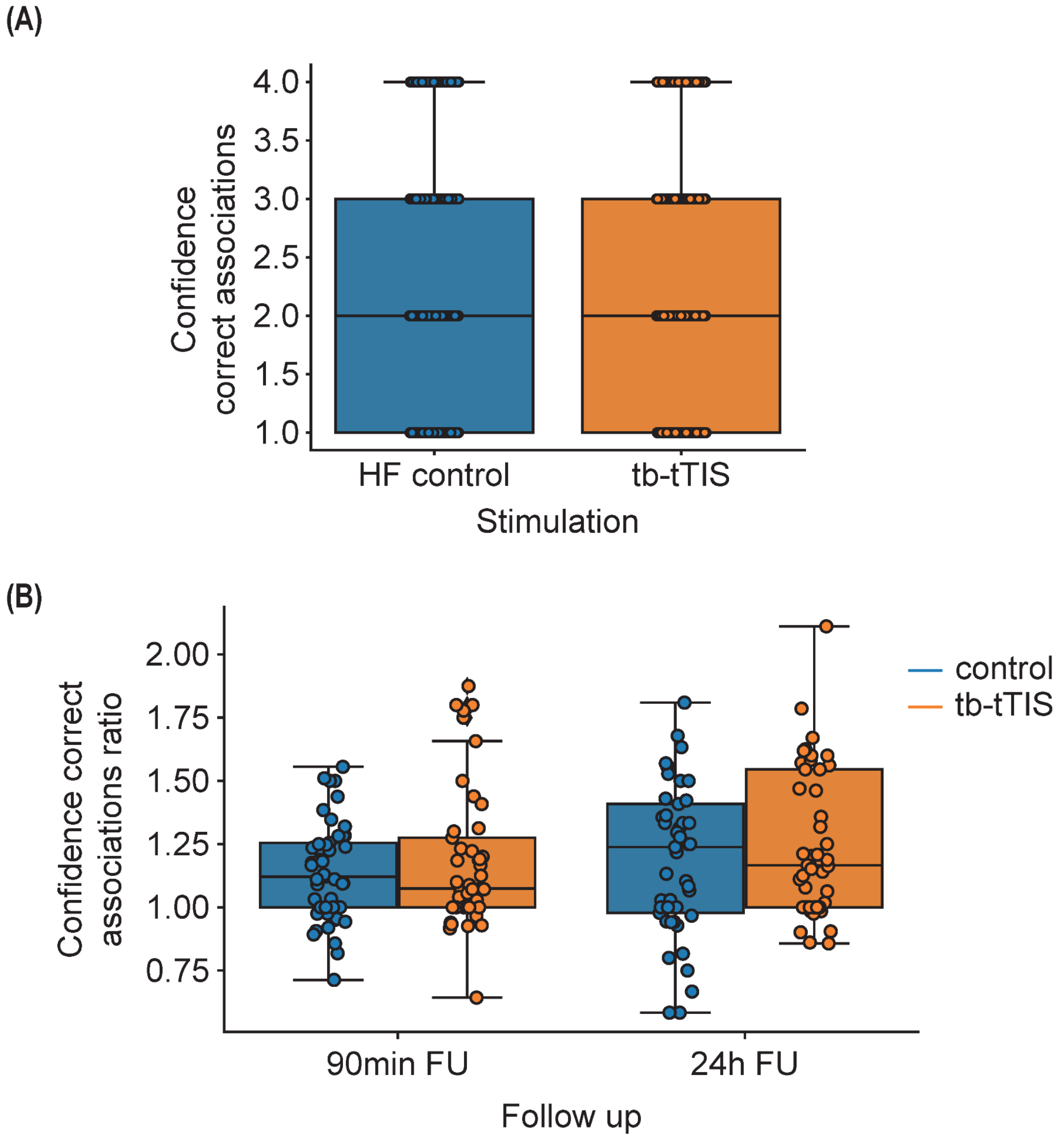


**Fig S3. Confidence results**

(A) Distribution of the confidence during stimulation blocks. No significant difference was found. (B) Confidence during the follow-up 90 minutes and 24 hours after stimulation with respect to the last stimulated block. No significant differences were found.

### Study participant selection criteria

**Exclusion criteria**

- Unable to consent
- Severe neuropsychiatric (e.g., major depression or severe dementia) or unstable systemic diseases (e.g., severe progressive and unstable cancer or life-threatening infectious diseases)
- Severe sensory or cognitive impairment or musculoskeletal dysfunctions prohibiting instruction comprehension or experimental task performance
- Inability to follow or noncompliance with the procedures of the study
- Contraindications for noninvasive brain stimulation (NIBS) or MRI:
    - Electronic or ferromagnetic medical implants/device; non-MRI compatible metal implant
    - History of seizures
    - Medications that significantly interact with NIBS are benzodiazepines, tricyclic antidepressants and antipsychotics
- Regular use of narcotic drugs
- Left-handedness
- Pregnancy
- Request to not be informed in case of incidental findings
- Concomitant participation in another trial involving neuronal plasticity probing.
